# Agentomics: An Agentic System that Autonomously Develops Novel State-of-the-art Solutions for Biomedical Machine Learning Tasks

**DOI:** 10.64898/2026.01.27.702049

**Authors:** Vlastimil Martinek, Andrea Gariboldi, Dimosthenis Tzimotoudis, Mark Galea, Elissavet Zacharopoulou, Aitor Alberdi Escudero, Edward Blake, David Čechák, Luke Cassar, Alessandro Balestrucci, Panagiotis Alexiou

## Abstract

**Motivation:** Extracting knowledge from biomedical data is crucial for advancing our understanding of biological systems and developing novel therapeutics. The quantity, quality, and resolution of biomedical data constantly evolves, requiring the automation of biomedical machine learning (ML). Existing Automated ML tools lack flexibility, while Large Language Models (LLMs) struggle to consistently deliver reproducible machine learning codebases, and existing LLM Agent-powered solutions lag behind human-engineered ML models.

**Results:** Here, we introduce Agentomics, an autonomous LLM-powered agentic system for end-to-end ML experimentation. Given a biomedical dataset, Agentomics implements various ML modeling strategies, and produces a ready-to-use ML model. Agentomics introduces strict validation checkpoints for standard ML development steps, allowing gradual development on top of working code with defined interfaces and validated artifacts. Further, it offers native support for biomedical foundation models that can be leveraged during experimentation. The generic nature of Agentomics allows the user to create ML solutions for a large variety of datasets and use various LLMs. We evaluate Agentomics across 20 datasets from the domains of Protein Engineering, Drug Discovery, and Regulatory Genomics. When benchmarked against other agentic systems, Agentomics outperformed them in all tested domains. When benchmarked against human expert solutions, Agentomics generated novel state-of-the-art models for 11/20 established benchmark datasets.

**Availability and Implementation:** Agentomics is implemented in Python. Source code and documentation are freely available at: https://github.com/BioGeMT/Agentomics-ML.

## Introduction

The past decades have witnessed the transformation of molecular biology into a truly data-driven science, in large part due to the growth in the quantity and variety of molecular biology data generated by technologies such as mass spectrometry and high-throughput sequencing (Marx, 2013; Stephens et al., 2015).

An important step in the analysis workflow for biomedical data is the use of Machine Learning (ML) techniques. Automated Machine Learning (AutoML) frameworks, such as Auto-sklearn (Feurer et al., 2015), AutoGluon (Erickson et al., 2020), TPOT (Olson and Moore, 2016), and H2O (LeDell et al., 2020), can automate the model selection and hyperparameter optimization steps of the ML development process. These frameworks are reliant on pre-defined data representations and limited architecture search (Azevedo et al., 2024). Thus, their utility is limited when applied to biomedical data, which is sometimes best represented in non-standard domain-aware representations. Additionally, new data modalities are often introduced, and slowly standardised. Accurate predictive models for biomedical data may require custom deep neural network architectures or the use of foundation models.

The capability of Large Language Models (LLMs) to generate code is already assisting general software engineering tasks. For ML experimentation, LLM-generated code can be manually executed in a run environment and incorporated into the ML pipeline. While this approach enables the development of customised data representations, as well as task-specific model architectures, it also introduces important limitations. In particular, the absence of an error-correction mechanism often results in low success rates for generated code, requiring human oversight for refinement and debugging (Jimenez et al., 2024). LLMs can be extended with tools that enable them to interact with an environment, and perform tasks such as code execution. Such implementations have come to be referred to as LLM Agents (Wang et al., 2024). Multi-step LLM Agents can also receive and react to error messages by adjusting their tool calls, effectively creating a loop that can be utilised for software development.

Recently, several specialised LLM Agents have been introduced that are able to autonomously develop ML models. These were categorised (Miller et al., 2025) as ‘generalist’ ML coding agents (AIDE (Jiang et al., 2025), MLAgentBench (Huang et al., 2024)), or more specialised ‘biomedical’ ML coding agents (STELLA (Jin et al., 2025), Biomni (Huang et al., 2025)).

AIDE (Jiang et al., 2025) uses a tree-search methodology, treating each potential ML solution as a node. It iteratively refines promising candidates, achieving top performance among Agent frameworks on benchmarks such as MLE-Bench (Chan et al., 2025).

MLAgentBench (Huang et al., 2024) uses the ReAct (Yao et al., 2023) and Reflexion (Shinn et al., 2023) prompting frameworks to guide an LLM-based Agent to construct ML solutions for problems from diverse domains (text, images, graphs, and tabular data).

STELLA (Jin et al., 2025) is a multi-agent system composed of manager, developer, critic, and tool creation agents. It improves results over time by gathering successful strategies in a template library. STELLA reports outperforming LLM methods and other Agents on various biomedical research tasks. Biomni (Huang et al., 2025) is a general-purpose biomedical Agent that uses an expert-vetted environment of pre-installed tools to assist with biomedical tasks, and retrieves tools from this set based on user goals. It beats various baselines on biomedical knowledge and reasoning, and demonstrates the capability to create multi-omics analysis workflows.

Despite their distinct approaches, LLM Agents have largely underperformed compared to human experts on a variety of biomedical ML benchmarks (Miller et al., 2025).

Beyond performance, LLM Agents that can autonomously execute code pose a security risk to the environment in which they operate. Implementing security measures to constrain LLM Agents adds overhead to the successful use of such methods. A lack of such measures can have catastrophic consequences, such as data loss due to execution of malicious code on the host machine.

Biomedical datasets are often sensitive and may be legally restricted in sharing or publishing. Support for local and open-source LLMs, that do not require API calls to a provider’s server, is important in order to securely use LLM Agents for ML on real biomedical data.

In this study, we present Agentomics, an autonomous LLM Agent for automating biomedical end-to-end ML experimentation. When benchmarked against other agentic systems, Agentomics outperformed them in all tested domains. When benchmarked against human expert solutions, Agentomics generated novel state-of-the-art models for 11/20 established benchmark datasets across the biomedical domains of Protein Engineering, Drug Discovery, and Regulatory Genomics.

Agentomics uses an iterative process to explore various ML strategies, while being able to use domain-specific foundation models to enhance its modeling capabilities. It also supports a wide variety of biomedical data modalities including, but not limited to, DNA, RNA, protein, and small molecule sequences. A single Agentomics run produces all necessary artifacts needed for running inference and retraining models with new data. Additionally, it produces a report summarising the final model and its predictive performance. All runs are fully containerised in a secure environment and prevent test set leakage and metric hallucinations. Agentomics supports a large selection of LLM providers and models, including local LLMs that can be used to securely retain sensitive biomedical data on the host machine.

## Methods

Agentomics is an autonomous system based on LLM Agents for end-to-end ML experimentation. Agentomics has been designed to produce ML models for classification and regression tasks. It ingests a single dataset, experiments with multiple strategies, and outputs a single ML model and all necessary artifacts for using it.

### Interface

The inputs of Agentomics are: **(a)** a training dataset file that will be available to agents, **(b)** an optional dataset description file that will be injected into the agent’s base prompt, **(c)** an optional test dataset file that will be used at the end of the run to report metrics to the user, and **(d)** an optional validation dataset file for early stopping and estimating performance, which, if not provided, is created by Agentomics by splitting the training dataset. All dataset files are required to be in a CSV format and contain a label column.

The outputs are: **(a)** scripts with a unified interface and the final model artifacts necessary for inference and re-training, **(b)** a conda environment necessary to run the produced scripts, **(c)** a PDF report including the summary of the chosen ML approach and train, test, and validation metrics, and **(d)** any miscellaneous files (e.g. exploratory scripts and results) that the LLM Agent created during the run.

Among the outputs, a successful run produces two scripts with a unified interface. The training script can be used to re-train the ML model, and always takes as input **(a)** a train data file in the same format as the original training file, and **(b)** a validation data file. It produces a new folder with all necessary artifacts for inference. These artifacts can then be passed to the second script, the inference script, to predict labels of new data. This takes as input a single file in the same format as the original training data, minus labels. It produces an output file with predicted labels for all samples in this input file.

The run can be further customised by various parameters, including the maximum time duration of the run, LLM backbone, and validation metric to use for optimisation. Additionally, the user can set a split budget, specifying for how long a run is allowed to re-split data into new train and validation sets, and a baseline budget, specifying how many initial iterations should implement only simple baseline ML models.

### Security and Privacy

Agentomics operates within an isolated Docker container with read-only mounts. This prevents any changes to the user’s host system outside of this Docker container. Agentomics optionally supports deployment with locally-hosted LLMs. This capability addresses privacy concerns in biomedical datasets.

### Tools

Agentomics uses six tools to enable agents to interact with their execution environment, which contains a pre-installed conda environment with packages like scikit-learn, torch, and transformers.

1. `The bash tool` enables running arbitrary bash commands and installing packages into the agent’s conda environment. It returns the console output of the command. This tool can create, edit, and remove files in the workspace.
2. `The python write tool` checks the syntax of a python file content and writes it into the Agentomics workspace. It returns whether the writing was successful. This tool can create files in the workspace.
3. `The python run tool` executes a python file with provided keyword arguments and returns the console output of the script and its run duration. This tool can create, edit, and remove files in the workspace.
4. `The edit tool` replaces specific content in a file and allows editing existing files with low token usage. It returns how many edits were done.
5. `The foundation models info tool` either returns a summary of all the available foundation models or detailed information and code snippets of a specific available foundation model.
6. `The final result tool` is called by the agent at the completion of each step to return the step’s structured output in JSON format. The output of each tool is capped at 5000 characters to prevent flooding of the LLM context. All tools can return error codes, messages, and error stack traces to the agent’s context.

### Foundation Models

Agentomics runs have access to several pre-downloaded foundation models from various biomedical domains. Six variants of ESM-2 (Lin et al., 2023) for protein sequences, five variants of HyenaDNA (Nguyen et al., 2023) and nine variants of NucleotideTransformer (Dalla-Torre et al., 2025) for DNA sequences, three variants of RiNALMo (Penić et al., 2025) for RNA sequences, and seven variants of ChemBERTa (Ahmad et al., 2022) and one variant of MolFormerXL (Ross et al., 2022) for small molecules.

An Agentomics run can retrieve the documentation and example code snippets of the available foundation models through the `foundation models info tool.`This allows generated python scripts to quickly implement architectures leveraging pre-trained foundation models. The list of available foundation models is easily extensible through a configuration file, allowing the addition of Hugging Face (Wolf et al., 2020) models.

### Predefined Steps

Agentomics follows a sequence of predefined steps (Figure 1). Each predefined step results in a structured output that serves as a concise summary for subsequent steps, reducing the overall context usage. Each step is implemented by a separate LLM Agent with a clean context that is prompted with **(a)** base prompt containing guidelines, **(b)** structured outputs of all previous steps, and **(c)** step-specific instructions. Each LLM Agent can use any of the tools from the Tools section and may use them repeatedly to interact with a shared environment (e.g. explore data, create files, execute code, observe results).

**Fig. 1.**
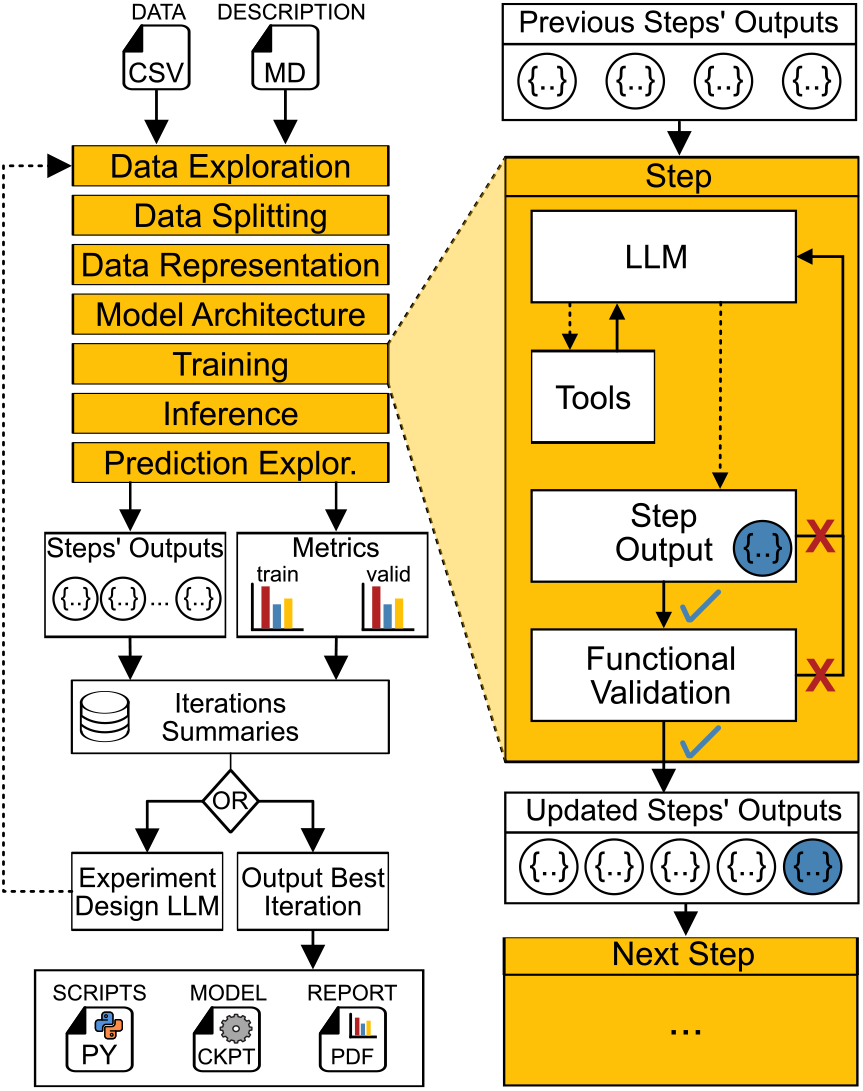
Architecture of Agentomics. Using input data and a description, Agentomics sequentially completes predefined ML experimentation steps using LLM Agents. Each step receives the outputs of previous steps, and is structurally and functionally validated, and potentially retried, before being considered successful. Once all steps are completed, their outputs and model metrics are saved as an Iteration Summary. The Experiment Design LLM then uses Iteration Summaries from all completed iterations to create instructions for each step of the next iteration. Once a run ends, Agentomics outputs files of the best iteration (based on the validation score), including training and inference scripts, model files, and a report including metrics and summarising the implemented strategy. Dotted arrows represent LLM-dependent actions.

Eventually, the Agent must submit the step structured output through the `final result tool.` All tool call messages are kept in the Agent’s context until the step is completed and the final step result has successfully passed validation.

Output must pass structured validation (valid JSON) and step-specific functional validation (Figure 1) before being considered completed. If validation fails, the LLM Agent receives error messages and must continue implementing the same step and re-submit the step output. This validation-retry loop continues until the validation passes, guaranteeing that subsequent steps operate on validated outputs and artifacts. If an Agent exceeds the retry limit, Agentomics restarts from the first step.

The steps can be summarized as follows:

1. *Data Exploration*: task is to explore data and output descriptive statistics, feature analysis, and domain-specific insights.
2. *Data Splitting*: task is to split the input data into training and validation sets and provide their paths. The following are programmatically validated: **(a)** split files existence and naming **(b)** retaining correct column names **(c)** non-overlapping samples between the splits. This step is skipped if the user provided a validation dataset.
3. *Data Representation*: task is to output text description of the intended data representation approach.
4. *Model Architecture*: task is to output text description of the intended model architecture and hyperparameters.
5. *Training*: task is to implement a training script and output its path, train a model using the split data, output a folder path containing training artifacts, and output a text summary of the implementation. The following are programmatically validated: **(a)** training script existence, accepted parameters, and naming **(b)** training artifacts existence, and naming **(c)** training script runability using a different subset of data, and production of the same artifacts as the submitted trained model.
6. *Inference*: the task is to implement an inference script, and output its path and a text summary of the implementation. The following are programmatically validated: **(a)** Inference script existence, accepted parameters, and naming **(b)** inference script runability using a different subset of data, and production of predictions for all samples in a correct format.
7. *Prediction Exploration*: task is to output statistics and insights from exploring model validation predictions and identifying potential prediction biases.

Once this last step is completed, the produced training artifacts and inference script are used to programmatically run inference on the validation and train splits. The predictions are then used to compute various metrics (Accuracy, AUROC etc) for both splits. This allows an objective comparison of training and validation metrics for different strategies without relying on LLM-reported metrics. These 7 steps are defined as an *iteration* of the run. Each successful iteration produces a single working ML model.

### Iterative Experimentation and Model Selection

At the end of each iteration, a model is produced (*M* in Line 6, Algorithm 1), which we use to generate predictions and obtain training and validation metrics (Line 12-13, Algorithm 1). These, together with structured outputs of all 7 steps (*s* in Line 6, Algorithm 1), create an iteration summary (Line 18, Algorithm 1).

At the beginning of a new iteration, we prompt the experiment design LLM to generate detailed instructions for each step (Line 4, Algorithm 1) that are used to extend the base prompt of that iteration (Line 5, Algorithm 1). The prompt for the experiment design LLM contains the iterations’ summaries of all completed iterations (Line 18, Algorithm 1), enabling it to make connections between each iteration’s ML strategy and its performance. Additionaly, it contains the dataset description, as well as information about the available GPU and CPU resources, remaining runtime, name of the validation metric to optimise, and information about available tools and foundation models. If configured, it also specifies whether the next iteration is allowed to split or re-split the data into training and validation sets, and whether the architectures should be limited to baseline models. In the prompt, we also include guidelines to reduce cases of agents over-optimising one strategy through incremental changes and instead focus on a more exploratory approach. If available for the given LLM backbone, this experiment design LLM call is set to high-reasoning effort.

An Agentomics run uses a time and/or an iteration limit. All iterations have access to the same file system and can repurpose models and scripts from previous iterations. After this limit is reached the run ends, and the final output falls back to the completed iteration with the best validation metric score (output iteration) (Line 20, Algorithm 1). Only iterations using the latest train-validation split are compared against each other to preserve fair comparisons (Lines 7-11, Algorithm 1). The specific validation metric to use for determining the best iteration can be adjusted by the user. If a test set is provided with the dataset, files and scripts from this iteration are then used to programmatically run inference and compute test metrics (Line 20, Algorithm 1). Note that the test set is completely isolated from any of the LLM Agents at all times in order to prevent any exploitation or data leakage.

#### Algorithm 1

Agentomics Iterative Experimentation

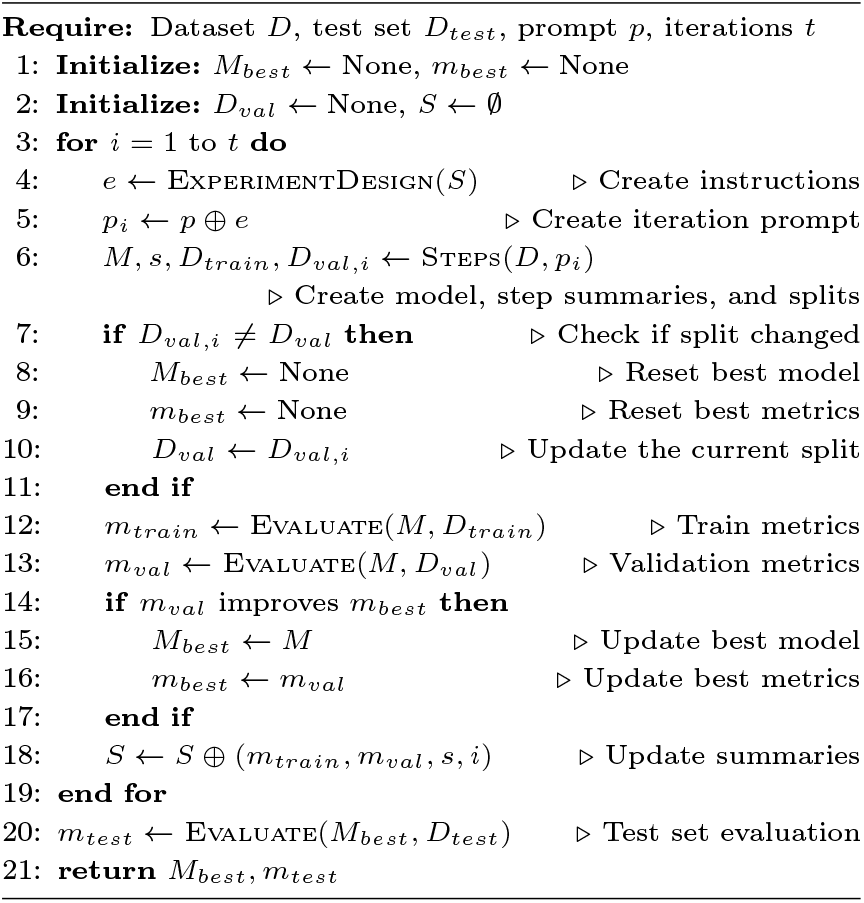

### Evaluation

To evaluate Agentomics in biomedical ML experimentation we use a set of standard and established benchmarks with existing state-of-the-art models, namely ProteinGym (Notin et al., 2023) for Protein Engineering tasks (N=6), Polaris Hub (Ash et al., 2025) for Drug Discovery tasks (N=9), and Genomic Benchmarks (Grešová et al., 2023) and miRBench (Sammut et al., 2025) for Regulatory Genomics tasks (N=5).

For Regulatory Genomics tasks, we compare against a leaderboard of expert human-engineered models, including the current state-of-the-art (Qiao et al., 2024; Yu et al., 2025; Thoutam and Ellsworth, 2024; Yang et al., 2025; Sammut et al., 2025). The leaderboard has been created by manually curating all published manuscripts that referenced the dataset in use, and reported test set metrics. The leaderboard is available as part of the Agentomics code repository. We also compare against a Zero-Shot LLM baseline that generates a solution in a single response. For Protein Engineering and Drug Discovery tasks we follow the BioML-bench framework (Miller et al., 2025), which enables us to compare against various autonomous LLM agents, as well as a leaderboard of best human-engineered solutions (Tables 1 and 2).

**Table 1.** Average performance of LLM agents against a leaderboard of expert human-engineered models. Completion % represents the proportion of runs that successfuly produced predictions. Leaderboard % is calculated only from successful runs.

| Domain | Agent Name | Leaderboard(%) | Completion(%) |
| --- | --- | --- | --- |
| Protein Engineering | Agentomics | **75.92 ± 33.96** | **100.0** |
| | AIDE | 35.99 ± 10.96 | 75.0 |
| | MLAgentBench | 13.52 ± 9.18 | **100.0** |
| | STELLA | 15.41 ± 6.81 | 87.5 |
| | Biomni | 29.93 ± 11.04 | **100.0** |
| Drug Discovery | Agentomics | **34.29 ± 27.67** | **100.0** |
| | AIDE | 31.06 ± 8.88 | 80.6 |
| | MLAgentBench | 22.45 ± 5.93 | **100.0** |
| | STELLA | 12.34 ± 6.62 | 91.7 |
| | Biomni | 32.55 ± 8.61 | **100.0** |
| Regulatory Genomics | Agentomics | **63.85 ± 39.56** | **100.0** |
| | Zero-Shot | 24.68 ± 25.83 | 80.0 |

**Table 2.** Comparison between Agentomics best-of-3 models and human best models across benchmark datasets. Descriptions of Protein Engineering and Drug Discovery datasets are adapted from BioML-bench (Miller et al., 2025).

| Dataset | Description | Dataset Size | Metric | Agentomics | Human best |
| --- | --- | --- | --- | --- | --- |
| Human Enhancers Cohn | Predict experimental human enhancers versus size-matched random genomic regions. | 27 791 | ACC | **0.759** | 0.747 |
| Human Enhancers Ensembl | Predict human enhancer elements against random genomic background sequences. | 154 842 | ACC | 0.916 | **0.933** |
| Human OCR Ensembl | Predict open chromatin regions against size-matched random genomic sequences. | 174 756 | ACC | **0.825** | **0.825** |
| Drosophila Enhancers Stark | Predict Drosophila melanogaster enhancer sequences against random genomic controls. | 6 914 | ACC | **0.838** | 0.772 |
| AGO2 CLASH Hejret2023 | Predict microRNA–target interactions using AGO2 CLASH-derived chimeric RNA pairs. | 8 193 | AUPRC | **0.880** | 0.860 |
| SPIKE-SARS2-Starr-2020 | Predict ACE2 binding fitness for SARS-CoV-2 spike RBD single-site variants. | 3 803 | $r_s$ | **0.934** | 0.747 |
| SPA-STAAU-Tsuboyama-2023 | Predict stability of *Staphylococcus aureus* protein A multi-substitution variants. | 1 036 | $r_s$ | **0.986** | 0.796 |
| PSAE-PICP2-indels | Predict stability effects of indel variants in photosystem I subunit IV. | 1 220 | $r_s$ | **0.954** | 0.847 |
| CBX4-HUMAN-multi-sub | Predict stability of human CBX4 SUMO ligase multi-substitution variants. | 918 | $r_s$ | **0.981** | 0.848 |
| Q8EG35-SHEON-indels | Predict organismal fitness for *Shewanella* MtrA protein indel variants. | 332 | $r_s$ | **0.949** | 0.775 |
| CSN4-MOUSE-indels | Predict stability of mouse COP9 signalosome subunit 4 indel variants. | 197 | $r_s$ | **0.979** | 0.906 |
| polaris-pkis2-egfr-wt-c-1 | Classify EGFR wild-type kinase inhibitors. | 496 | AUPRC | 0.752 | **0.866** |
| polaris-adme-fang-hclint-1 | Predict human liver microsomal intrinsic clearance from SMILES. | 2 229 | $r_p$ | 0.697 | **0.778** |
| polaris-adme-fang-hppb-1 | Predict fraction unbound in human plasma protein binding. | 126 | $r_p$ | 0.815 | **0.888** |
| polaris-adme-fang-solu-1 | Predict compound solubility from SMILES using standardised ADME protocols. | 1 578 | $r_p$ | 0.628 | **0.770** |
| tdcommons-cyp2d6-substrate | Classify CYP2D6 enzyme substrates for drug–drug interaction risk. | 532 | AUPRC | 0.690 | **0.724** |
| tdcommons-lipophilicity | Predict lipophilicity (logP) from SMILES. | 3 360 | MAE | 0.516 | **0.392** |
| tdcommons-herg | Classify hERG potassium channel blockers to assess cardiotoxicity risk. | 523 | AUROC | **0.892** | 0.891 |
| tdcommons-bbb-martins | Classify blood–brain barrier penetration. | 1 624 | AUROC | 0.892 | **0.933** |
| tdcommons-caco2-wang | Predict intestinal permeability in Caco-2 cell assay. | 728 | MAE | **0.277** | 0.282 |

For each dataset, we run Agentomics and generate a ML model, followed by computing metrics using a held-out test set. For Protein Engineering and Drug Discovery datasets, we use the BioML-bench framework to run Agentomics and follow the exact same evaluation criteria and time restrictions as competing agents (Miller et al., 2025). Agents are compared based on the percent of human leaderboard solutions they outperform.

For each dataset, we run 3 replicates for a duration of 8 hours each, on a Linux machine with access to NVIDIA RTX A4000 GPU and 48 CPU cores. For all experiments, we ensure the Agents do not have access to the test portions of datasets. We used OpenAI GPT-5.1-Codex-Max as the LLM backbone for all Agentomics and Zero-Shot runs. The split budget, during which the Agent is allowed to re-split train and validation data, was set to half of the run (4 hours). The baseline budget, during which the Agent is prompted to use only simple architectures and data representations to set a baseline, was set to 4 iterations. The validation metrics were dependent on the dataset, based on which metric is reported by state-of-the-art models or BioML-bench benchmarks. Classification datasets used either Accuracy (ACC), area under the precision-recall curve (AUPRC), or area under the ROC curve (AUROC). Regression datasets used either mean average error (MAE), Pearson correlation coefficient (*r*_*p*_), or Spearman correlation coefficient (*r*_*s*_).

A complete trace of all Agent actions and code produced during the run is exported. We calculate train, validation, and test metrics, as well as money spent on LLM API calls for each run. We also measure success rate: the percentage of runs that **(a)** produce predictions, **(b)** produce a valid conda environment, **(c)** produce an inference script that can be used to infer arbitrary data of the same structure, and **(d)** produce a training script that can be used to re-train the model with arbitrary data of the same structure.

## Results

### Agentomics outperforms other agents at Protein Engineering and Drug Discovery

In order to evaluate the performance of Agentomics against other LLM Agents, we evaluated our method based on the percentage of human leaderboard solutions it outperforms, following the methodology of the BioML-bench (Miller et al., 2025) framework, on Protein Engineering (N=6) and Drug Discovery (N=9) tasks. We performed triplicate runs on each dataset, and compared directly against leaderboard percentile of successful runs of four other LLM Agentic methods as reported by BioML-bench (Table 1).

Agentomics achieved mean percentiles of 75.92, 34.29, and 63.85 for Protein Engineering, Drug Discovery, and Regulatory Genomics respectively, outperforming all other benchmarked methods in all domains.

### Agentomics produces new state-of-the-art for the majority of tested datasets

We have evaluated the best replicate (out of 3) for Agentomics against the reported human state-of-the-art on each of the 20 benchmark datasets. For Protein Engineering tasks, Agentomics outperformed human state-of-the-art in all 6/6 datasets. In Drug Discovery tasks it achieved new state-of-the-art in 2/9 datasets. In Regulatory Genomics, it matched or improved on the current human state-of-the-art in 4/5 datasets (Table 2).

### Agentomics indirectly optimises for generalisation

We theorised that optimising validation performance across iterations may result in indirect overfitting to the validation data and thus hurt generalisation. To estimate if Agentomics optimises generalisation, we computed the Pearson correlation coefficient of the directly optimised validation score against the hidden test score over all iterations of all runs (Figure 2A). We observed high validation-test score correlations in the Regulatory Genomics runs (mean 0.986 ± 0.026), and slightly lower correlations in the Protein Engineering runs (mean 0.746 ± 0.226), and Drug Discovery runs (mean 0.807 ± 0.286). Validation-test score correlation across all runs was generally high (Figure 2) with a median value of 0.964. This suggests that optimising validation performance generally does not overfit to the validation data, and translates well to unseen test set performance. This finding highlights the ability of Agentomics to autonomously split train-validation data in a representative manner that optimises well for generalisation.

**Fig. 2.**
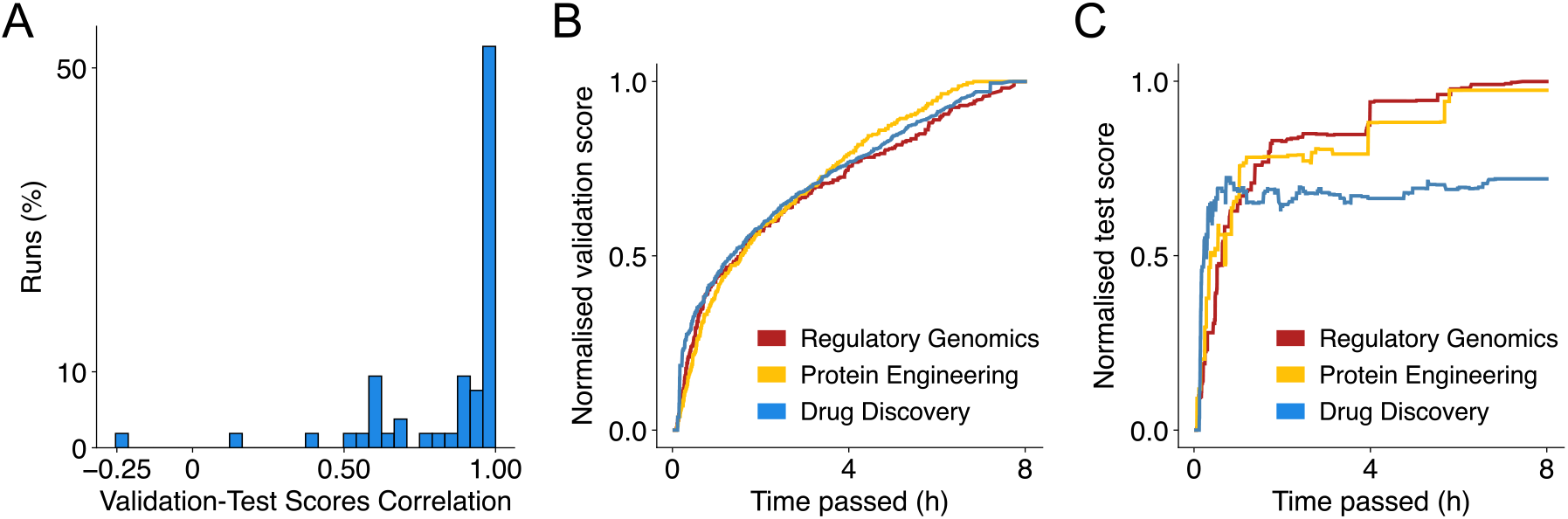
Agentomics optimises for generalisation. (A) Distribution of Validation-Test Scores Pearson Correlations from all runs. (B) Normalised validation score and (C) Normalised test score (score on the test set if the run stopped at that time point) over 8 hours averaged from all runs stratified by domain. Value of zero represents the worst model of each run and a value of one represents the best model based on validation and test scores respectively.

### Agentomics discovers better performing ML strategies over time

A single Agentomics run implements various ML strategies across many iterations. If an iteration’s submitted model achieves a new best validation score, that iteration is set to be the output iteration and will be the result of the finished run. We computed the validation and test scores for a would-be output iteration across the duration of a run, normalised them per run (0 meaning lowest score from any iteration, 1 meaning highest score from any iteration), and plotted their averages stratified by domain (Figure 2B,C).

In all domains, we observe that Agentomics continues to improve the validation score over time consistently. For Regulatory Genomics and Protein Engineering tasks, we observed that test scores also improved over time, although in a more staggered manner. However, Drug Discovery runs stagnated their test performance early, around the 1 hour mark. Drug Discovery output iterations also had lower normalised test scores on average, pointing to a phenomenon where the best test scores were reached in iterations with the non-best validation scores, and therefore not being selected as output iterations.

We do not observe a substantial dip in the normalised test score over time for any group, implying that on average the test scores were not negatively influenced by continuing the run past the point when test score stopped improving (e.g. due to overfitting to the validation set).

### Agentomics explores a wide variety of strategies

To investigate the diversity of solutions explored during Agentomics runs, we analysed the architectures implemented during the best-of-3 run for each of the 20 benchmarked datasets and plot their distribution in all iterations (N=364, red bars) and just the output iterations (N=20, blue bars) (Figure 3). We observe that Agentomics explores a wide variety of strategies across its runs, implementing both traditional architectures like linear regression or support vector machines, as well as deep learning architectures like transformers and convolutional neural networks. The most common approach was to use ensemble methods like XGBoost, random forests, and custom ensembles composed of various individual architectures. We also observe that Agentomics commonly utilises foundation model embeddings, using them in 20% of output iterations.

**Fig. 3.**
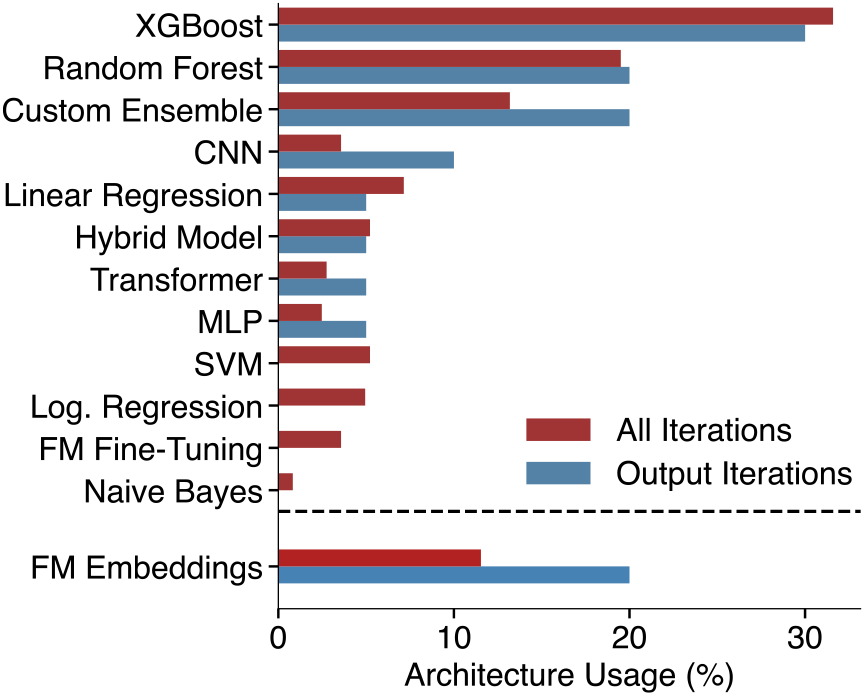
Agentomics implements varied strategies. Distribution of architecture types implemented by the best Agentomics run for each dataset, shown separately for all iterations (N=364, red bars) and output (best validation score) iterations (N=20, blue bars). Additionaly, a percentage of iterations that used foundation model embeddings is shown. Custom Ensemble architecture represents non-XGBoost, non-Random Forest, ensemble methods composed of various other architectures. Hybrid model architecture represents non-ensemble architectures that combine two or more simple architectures. FM=Foundation Model.

### Case Study: AGO2 CLASH Hejret 2023

We further analyse how Agentomics tackles the AGO2-CLASH Hejret 2023 benchmark, comprising 8,193 miRNA–target chimeric pairs identified via AGO2 CLASH technology (Hejret et al., 2023). Each sample consists of two RNA sequences, and the task is to decide whether they bind or not in the context of the AGO2 protein. This dataset has been extensively studied using a range of machine learning architectures (Sammut et al., 2025). As a case study, we investigate the best-performing 8-hour Agentomics run, which outperformed the human state-of-the-art (AUPRC 0.880 vs. 0.860).

The run began with a logistic regression baseline on one-hot encoded RNA sequences, followed by gradient-boosted trees, and standard deep learning approaches including CNNs and transformers. The following iterations implemented interaction matrices encoding Watson–Crick and wobble base pairing, a representation closely aligned with known miRNA–target biology and similar to that used in expert-designed models. Notably, these representations and models were not provided to the agent. Agentomics further fine-tuned a foundation model (Nucleotide Transformer *∼* 50M parameters (Dalla-Torre et al., 2025)), experimented with cross-attention dual encoders, and implemented CNN ensembles of residual, depthwise-separable, and ConvNeXt-style Liu et al. (2022) architectures. Multiple ensemble strategies were explored, including weighted averaging, stacking with logistic regression and XGBoost, and large ensembles of prior high-performing solutions. The output solution was a probability-weighted combination of five previously implemented ensembles, explored in five different iterations, and achieved the strongest overall test set performance.

Overall, this case study demonstrates that Agentomics iteratively explores, refines, and recombines a wide range of modeling strategies, ultimately exceeding expert-designed solutions through extensive exploration of the strategy space.

### Agentomics is reliable, fast, and cost effective

LLM-based systems need LLM completions to generate tool calls and responses. All Agentomics experiments were run with the recommended GPT-5.1-Codex-Max backbone, which currently (Jan 2026) bills at 1.25$/M input tokens and 10$/M output tokens. We have tracked the spending of each of Agentomics runs, and find that 8-hour runs on average spend 9.4 ± 5.0 USD. The average cost of a single iteration is 0.45 ± 0.09 USD. The average duration for a single iteration is roughly 30 minutes, at 0.51 ± 0.32 hours. Over a total of 60 runs, Agentomics generated a working reusable solution every time, achieving a success rate of 100%.

### Agentomics is publicly available and open-source

We provide a user-friendly way to use Agentomics through a single command that will install necessary dependencies and guide user through configuring a run. Agentomics is open-source and can be used with a wide range of datasets and without requiring the user to have biomedical domain knowledge, programming experience, or machine learning expertise. Agentomics allows the user to customise each run through various parameters like maximum duration and LLM backbone.

## Discussion

Agentomics is an autonomous LLM Agent-based system for end-to-end biomedical ML experimentation that takes a dataset as input, iteratively designs and implements complete ML solutions, and returns validated and ready-to-use training and inference code as well as model artifacts. We evaluated Agentomics across three biomedical domains: Regulatory Genomics, Protein Engineering, and Drug Discovery, on a total of 20 established benchmark datasets. We found that Agentomics outperformed all other LLM Agents on the two domains where agent baselines were available (Protein Engineering and Drug Discovery), while also surpassing the best reported human engineered state-of-the-art in 11/20 datasets. The gains were pronounced in Protein Engineering, where Agentomics exceeded human leaderboard performance in 6/6 tasks, and in Regulatory Genomics, where it improved on human state-of-the-art in 4/5 tasks, showing that the effect is not confined to a single modality or benchmark suite. These results provide evidence that fully autonomous agentic ML systems can now surpass human expert ML models at scale across multiple biomedical domains. This shifts the practical frontier from LLM-assisted development to autonomous discovery of expert-level models.

There are many ways to implement LLM Agent-based systems for ML. Differences in agent architecture, tool access, and evaluation design can influence the effectiveness of such systems. Agents operating without isolated environments are susceptible to unsafe code execution and test label leakage, while unstructured agent architectures might hallucinate metrics and produce non-reusable outputs.

To guide the development of future LLM Agentic systems for robust ML, we would like to propose a set of standards: **(a)** runs should execute in a sandboxed environment (e.g. containerised with restricted mounts) to protect the user’s machine and data; **(b)** the test set must be strictly hidden from the agent to prevent data leakage; **(c)** metrics should be computed by deterministic evaluation code, not by the agent; and **(d)** to ensure reproducibility and transparency, the system should output reusable artifacts and scripts that can be inspected and used for training reproducibility and inference on arbitrary data.

Agentomics creates a validation set out of the original data and optimises its performance over multiple iterations. We observe that optimising this objective translates into improved generalisation on the hidden test set in most cases, as evidenced by high validation-test scores correlation (Figure 2A). Notable exception to this are Drug Discovery tasks, where test set metrics stagnate early and lag behind validation metrics.

Examining the best runs across datasets shows that there is no single predominant architecture for biomedical ML. Strong solutions were discovered through iterative experimentation, spanned different model families, and often leveraged biomedical foundation models. This positions AutoML methods that only explore limited architecture and representation spaces as not optimal for biomedical ML.

A limitation of our study is that it does not explore long running (8+ hours) experiments. Based on our results, certain datasets, where metric curves did not plateau, would likely have benefited from longer experimentation. Additionally, our experiments are limited to a single LLM backbone, and used limited hardware resources. Access to more GPU resources would shorten model training times and allow experimentation with larger models and datasets within reasonable timeframes. Finally, our experiments do not isolate individual components of the Agentomics architecture to measure their contribution through ablation study.

Analysing data from all runs, we observe high performance variance. While best replicates and average performance are strong, occasional runs produce relatively weaker models. Reducing this variance will be important for improving the reliability of Agentomics. Additionaly, Agentomics currently only supports data in CSV format, limiting its use cases for modalities such as images or specialised data formats.

Although motivated by biomedical tasks, Agentomics implements supervised ML models in a generic way as a sequence of validated steps. Additionaly, it is easily configurable and extendable with foundation models from any domain. This positions Agentomics as domain-agnostic and fit to be applied to diverse datasets.

## Conclusions

Systems like Agentomics point to a near future where autonomous ML experimentation becomes routine. They are increasingly affordable to run, fast enough for practical use cases, and robust enough to produce reusable and reproducible artifacts. As these agentic systems become more capable and widely adopted, the key risk is not that they will fail to produce models, but that poorly designed implementations will flood the field with irreproducible code and inflated metrics due to test set leakage, ultimately eroding trust in computational biomedical ML. With Agentomics, we demonstrate that Agentic system for ML can be developed in a way that mitigates these risks, guaranteeing reliability, security, and standardised interfaces needed to develop high quality scientific ML models at scale.

## Acknowledgments

VM, AG, DT, AB, EZ, and PA were supported by HORIZON-WIDERA-2022 BioGeMT (ID: 101086768). EB and LC were supported by HORIZON-EIC-2022-Pathfinderchallenges-01-03 TargetMI (ID:101114924), AAE, and PA were supported by University of Malta’s Data Integrity and Stewardship Cluster (DISC). DC was supported by the Bioinformatics Core Facility of CEITEC Masaryk University via the NCMG Research Infrastructure (LM2023067 funded by MEYS CR).

